# DeepToA: An Ensemble Deep-Learning Approach to Predicting the Theater of Activity of a Microbiome

**DOI:** 10.1101/2022.04.04.486969

**Authors:** Wenhuan Zeng, Anupam Gautam, Daniel H. Huson

## Abstract

**Motivation:** Metagenomics is the study of microbiomes using DNA sequencing. A microbiome consists of an assemblage of microbes that is associated with a “theater of activity” (ToA). To what degree does the taxonomic and functional content of the former depend on the (details of the) latter? More technically, given a taxonomic and/or functional profile estimated from metagenomic sequencing data, how to predict the associated ToA? Here we present a deep learning approach to this question. We use both taxonomic and functional profiles as input. We apply node2vec to embed hierarchical taxonomic profiles into numerical vectors. We then perform dimension reduction using clustering, to address the sparseness of the taxonomic data and thus make it more amenable to deep learning algorithms. Functional features are combined with textual descriptions of protein families or domains. We present an ensemble deep-learning framework DeepToA for predicting the ‘theater of activity” of microbial community, based on taxonomic and functional profiles. We use SHAP (SHapley Additive exPlanations) values to determine which taxonomic and functional features are important for the prediction.

**Results:** Based on 7,560 metagenomic profiles downloaded from MGnify, classified into ten different theaters of activity, we demonstrate that DeepToA has an accuracy of 98.61%. We show that adding textual information to functional features increases the accuracy.

**Availability:** Our approach is available at http://ab.inf.uni-tuebingen.de/software/deeptoa.

**Supplementary information:** Supplementary data are available at *Bioinformatics* online.

## 1 Introduction

Deep-learning algorithms are known to perform well on a wide range of problems coming from different disciplines of science (Jumper *et al*., 2021; Reichstein *et al*., 2019; Ardila *et al*., 2019; Bukhari *et al*., 2020). To achieve good performance on new problems, the data must satisfy certain requires (Najafabadi *et al*., 2015), and it is usually necessary to perform corresponding adjustments when applying deep-learning algorithms to a new domain.

Several deep-learning approaches have been developed to address microbiome-related questions, such as disease prediction (Oh and Zhang, 2020; Sharma and Xu, 2021), the annotation of antibiotic resistance genes (ARGs) (Li *et al*., 2021), microbial source tracking (Shenhav *et al*., 2019; Wu *et al*., 2021), and microbial community prediction (Thompson *et al*., 2019; Zha *et al*., 2020).

A microbiome consists of a collection of microbes that live in a specific “theater of activity” or ecological context (Whipps JM, 1988), and the total genomic information of the microbes is known as the metagenome of the microbiome (Handelsman, 2004). Analysis usually involves estimating the taxonomic and functional content of microbiomes, either from amplicon sequencing data (e.g. 16S rRNA sequences, (Caporaso *et al*., 2010)), or from metagenomic sequencing data (e.g. (Huson *et al*., 2016; Mitchell *et al*., 2020)).

The taxonomic and functional content of a microbiome is shaped, to a degree, by its theater of activity (The Human Microbiome Project Consortium, 2012; Berg *et al*., 2020). How strong is the influence? In particular, can one accurately predict the theater of activity from the taxonomic and functional profile of a sample? Here, we address this question using a deep-learning approach. We have trained and validated our approach using 7,560 metagenomic datasets downloaded from MGnify (Mitchell *et al*., 2020), classified into ten different theaters of activity, namely *Animal Digestive System, Food Production, Freshwater, Human Respiratory System, Human Skin, Mammal Gastrointestinal Tract, Marine, Plants, Soil* and *Wastewater*. For a given query sample, represented by a taxonomic and/or functional profile, our classifier returns a probability of membership for each of the ten classes.

In related work, the Earth Microbiome Project uses random forests to determine environmental factors (Smith *et al*., 2010). SourcePredict (Borry, 2019) uses dimension reduction followed by a KNN algorithm to classify and predict the origin of metagenomics samples. In a recent study (Wu *et al*., 2021), several machine learning algorithms are used to predict the dominant source of microbial contamination by using environmental and geographical data. The ONN4MST method uses an ONN (ontology-aware neural network) to embed biome ontology information into a hierarchical structure so as to improve performance of community-based microbial source tracking (Zha *et al*., 2020).

Some machine-learning approaches for predicting the theater of activity are based on taxonomic profiles of 16S rRNA sequencing data (Knights *et al*., 2011), which have limited taxonomic resolution and cannot assess functional content. Here we present an ensemble deeplearning framework DeepToA that is specifically designed for the analysis of taxonomic and functional profiles obtained from metagenomic data (see Figure 1). We provide the ToA prediction of metagenomics sample at our software server.

**Fig. 1.**
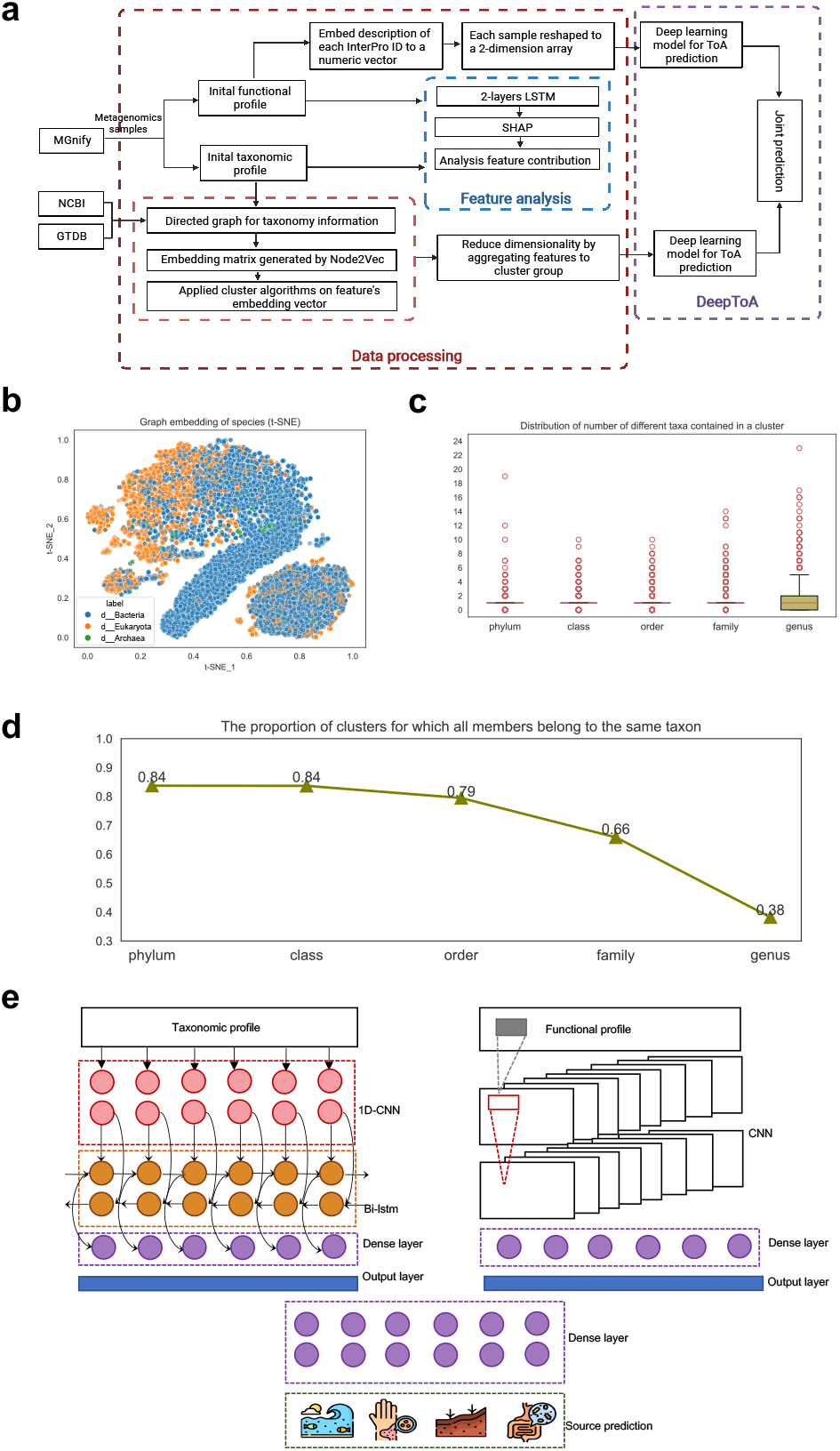
a Workflow for data collection, feature engineering and model building. Metagenomic samples are downloaded from MGnify and their taxonomic and functional profiles are extracted. Taxonomic and functional profiles are processed in the lower and upper part of the workflow, respectively. In the middle of the workflow, feature importance is assessed using SHAP values. Joint prediction using DeepToA is performed on the righthand side. b A t-SNE plot of species’ embedding vectors, colored by taxonomic domain. c The distribution of the number of different taxa that are contained in a cluster, for each taxonomic rank. The mean value is 1 for all ranks. d The proportion of clusters that are pure for a given taxonomic rank, that is, for which all members of the cluster belong to the same taxon of that rank. e Structure of the DeepToA model, with the taxonomic model left, the functional model right, and the combining and prediction layers at the bottom.

To train and evaluate DeepToA, we downloaded 7,560 metagenomic samples from MGnify (Mitchell *et al*., 2020). Each sample possesses a taxonomic profile based on the NCBI taxonomy (Schoch *et al*., 2020), a functional profile based on InterPro (Blum *et al*., 2021), and a known theater of activity. As described further below, we enhanced the samples by considering additional textual descriptions of the samples.

Taxonomic and functional profiles are usually represented by count tables, and for any given sample, the vast majority of entries will be zero. To address this, dimensionality reduction is required (Oudah and Henschel, 2018; Zhou *et al*., 2021; Sharma *et al*., 2020).

The first step in our approach is to convert taxonomic names lineage into numerical vectors. Popular pre-trained language models (Devlin *et al*., 2018; Peters *et al*., 2018) are not applicable here. Instead, we use both the NCBI taxonomy (Schoch *et al*., 2020) and the GTDB taxonomy (Parks *et al*., 2021) as a taxonomic reference tree for archaea, bacteria, eukaryota, and viruses, and extend this tree by attaching taxonomic lineages appeared in our taxonomic profile but not included in the reference tree yet. We apply node2vec (Grover and Leskovec, 2016) to this reference tree so as to obtain an embedding vector for each taxon.

We then calculate euclidean distances between the embedding vectors and use the AGNES algorithm to cluster them (which showed the best performance, as described below). The resulting clusters, which we will refer to as *processed taxonomic profiles*, are then used as input to a taxonomy-based deep-learning model for ToA prediction. Below, we show that the clusters reflect taxonomic relationships.

The second step in our approach is to process functional profiles. These are initially given as InterPro count tables, which we expand into three-dimensional tables by considering each features’ textual description. The resulting *processed tables* are used as input to a function-based deep-learning model for ToA prediction.

The taxonomy-based and function-based models are then combined into an ensemble deep-learning framework DeepToA, which achieves an accuracy of 98.61%.

To address explainable machine learning, we use a Bi-LSTM (bidirectional long short-term memory) model (Hochreiter and Schmidhuber, 1997) on the input taxonomic and functional profiles, respectively, and then compute SHAP (SHapley Additive exPlanations) values (Lundberg and Lee, 2017) to estimate feature importance, a widely-used approach for “explaining” deep-learning models (Rajpurkar *et al*., 2021; Arcadu *et al*., 2019; Yap *et al*., 2021). This will help to identify key taxa and functions associated with specific environmental conditions.

## 2 Materials and methods

### 2.1 Dataset download and preparation

We downloaded 7,560 metagenomic samples from MGnify (Mitchell *et al*., 2020). In more detail, in a first step, for each sample, we downloaded a taxonomic count table using the MGnify API. We used the QIIME script `merge_otu_tables.py` (Caporaso *et al*., 2010) to merge all tables into a single, initial table, containing 7, 560 rows (samples) and 117, 727 columns (taxa). Using the dimensionality reduction procedure described below, we clustered all taxa into 10, 000 classes and this reduced the taxonomy table to a “processed” table with 7, 560 rows (samples) and 10, 000 columns (taxonomic clusters).

In a second step, for each sample, we downloaded a functional count table using the MGnify API. That data was combined into a single initial table, containing 7, 560 rows (samples) and 13, 041 columns (InterPro IDs). We also downloaded the textual descriptions associated with the InterPro IDs from MGnify and embedded each description into a numerical vector of length 10 using doc2vec, as described below. This gives rise to a three-dimensional functional “processed” table with 7, 560 rows (samples), 13, 041 columns (InterPro IDs) and 10 additional values (the InterPro embedding vectors).

In this study, we use both the initial and processed datasets.

### 2.2 Computation of the taxonomy embedding matrix

#### 2.2.1 Tree structure

We downloaded taxonomic details on 4, 316 archaea and 254, 090 bacteria from the GTDB database release 202 (Parks *et al*., 2021), which is based on 258, 406 genomes organized into 47, 894 species groups. We also downloaded the taxonomy details on NCBI taxonomy database (Schoch *et al*., 2020), which covering 514 archea, 5, 294 bacteria, 64, 462 eukaryota and 1, 838 viruses.

All taxa mentioned in taxonomic profiles downloaded from Mgnify either included in the reference tree, or were added to the reference tree, together with the corresponding high-order taxa, if necessary. In result, we obtain an extended taxonomy that is based on both the GTDB taxonomy and the NCBI taxonomy, and is organized in eight usual taxonomic ranks Domain, Kingdom, Phylum, Class, Order, Family, Genus, and Species. We implemented the taxonomy as a rooted, directed tree, using the Networkx Python package (see https://networkx.org).

#### 2.2.2 Graph embedding

We ran the node2vec algorithm (Grover and Leskovec, 2016) on the taxonomic tree. In more detail, using 300 random walks per node and 20 nodes in each walk, we mapped each taxon *t* onto a 10-dimensional embedding vector *v*(*t*). The embedding vectors for all species are show in a t-SNE plot (van der Maaten and Hinton, 2008) in Figure 1b, indicating some clustering by domain.

We reshape the data by assigning to each taxon *t* a lineage-based vector of length 80 that is obtained as the concatenation *V* (*t*) = *v*(*t*_1_) ⊕ … ⊕ *v*(*t*_8_) of all embedding vectors *v*(*t*_1_), … (*t*_8_) of the taxa that lie on the path *t*_1_, *t*_2_, … *t*_8_ from the root of the taxonomy to the taxon *t*, one for each taxonomic rank, filling in missing data with zeros.

In total, we obtained 117, 727 different 80-dimensional vectors representing taxa.

### 2.3 Dimension reduction

To reduce the number of different taxonomic features to a more manageable number that is similar to the number of available samples, 7, 560, we decided to cluster the taxonomic features into 10, 000 clusters. To address this, we computed euclidean distances between all vectors and then evaluated the performance of different distance-based clustering algorithms, see Table 1.

**Table 1.** For eight different methods considered for clustering taxonomic vectors, we report the Calinski-Harabasz score, Silhouette score, Davies-Bouldin index, and total wall-clock time in minutes.

|  | Calinski-Harabasz | Silhouette | Davies-Bouldin | Time [m] |
| --- | --- | --- | --- | --- |
| K-means | 4,567.42 | 0.83 | 0.47 | 93.6 |
| DBSCAN | 61.16 | 0.75 | 1.01 | 0.83 |
| GMM | 4,548.56 | 0.83 | 0.46 | 464.62 |
| Spectral | 5.89 | -0.43 | 0.78 | 1,690.18 |
| Birch | 1,878.28 | 0.77 | 0.41 | 0.88 |
| AGNES | 4,978.53 | 0.85 | 0.48 | 26.39 |
| Mini-batch K-means | 1,005.92 | 0.79 | 1.02 | 8.48 |
| OPTICS | 114.08 | 0.84 | 1.05 | 50.62 |

We used three metrics to evaluate the performance of the clustering techniques, namely the Calinski-Harabasz score (Caliński and Harabasz, 1974), the Silhouette score (Rousseeuw, 1987), and the Davies-Bouldin index (Davies and Bouldin, 1979). In addition, we also compared running time.

Based on the results reported in Table 1, we selected the AGNES clustering algorithm for use in this study, as it has high Calinski-Harabasz score, the highest Silhouette score and low Davies-Bouldin score, suggesting more dense and well-separated clusters. The clustering gives rise to *r* = 10, 000 groups of taxa, which we denote by *C*_1_, …, *C*_10,000_.

After this preprocessing, any given taxonomic profile *t* associated with a metagenomic dataset can be represented as a vector *D* = (*d*_1_, …, *d*_*r*_) of length *r* = 10, 000, where 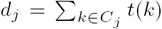 is the sum of counts over all taxa *k* that lie in cluster *C*_*j*_.

Let *T*_*m,n*_ = {*t*_*ij*_} denote the matrix of all original input taxonomic profiles, with *m* the number of samples and *n* the number of taxa, here *m* = 7, 560 and *n* = 117, 727. We will use *D*_*m,r*_ = {*d*_*ij*_} to denote the matrix of all “processed” taxonomic profiles, where *r* is the number of clusters, and, for every sample *i* and cluster *C*_*j*_, the entry 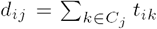 is the sum of counts over all taxa *k* of sample *i* that lie in cluster *C*_*j*_.

The GTDB and NBCI taxonomies are based on evolutionary relationships and it is important that the clustering of taxa described here reflects these relationships. This appears to be the case. In Figure 1d, for each higher taxonomic rank, we report the proportion of clusters that are pure in the sense that all members of the cluster belong to the same taxon of the given rank. We see that a large proportion of clusters are pure at the rank of Phylum (84%) and this drops to 38% at the rank of Genus, as is to be expected. In Figure 1c we show the distribution of number of different taxa that are contained in a cluster, for all higher taxonomic ranks. The mean count is 1 for all ranks.

### 2.4 Machine learning

We now describe the architecture of the neural network used for “theater of activity” prediction. We then discuss how to determine feature importance.

#### 2.4.1 Main neural network architecture in classification task

##### Taxonomic model

The processed taxonomic input data is represented as a table of counts, with rows representing samples and columns representing clusters of taxa. This is provided as input to a stacked combination of a 1D-CNN model (one-dimensional convolutional neural network) and an LSTM (long short-term memory) model. While a 1D-CNN architecture is usually used for text data and 1D signal data, an LSTM is specifically designed for processing long textual data. The combination of these two structures performs better than either model separately. We performed all training on the open-source framework Tensorflow-GPU Keras 2.6.0 (see https://www.tensorflow.org/). and we provide more implementation details further below.

##### Functional model

The original functional input data is provided as a table of counts, with rows representing samples and columns representing (13, 041) InterPro families. For each such family, we computed a 10-dimensional embedding vector of the textual description of the family, using doc2vec (Le and Mikolov, 2014), and thus obtained a two-dimensional 13, 041 *×* 10 matrix, which can be interpreted as a gray-scale image. This gives rise to the “processed” functional data, to we which we apply a two-layers CNN (to capture hidden rules), multiple max-pooling and dense layers, and an activation function (in the usual way).

##### Ensemble deep learning

In DeepToA, the taxonomic and functional deep learning models are combined into an ensemble model to perform “theater of activity” prediction together, as shown in Figure 1c.

#### 2.4.2 Explainable deep-learning prediction

For a given prediction of “theater of activity”, we would like to know which features play a role in the prediction. To address this, we designed a multi-categorical classification model that operates directly on the initial taxonomic and functional profiles.

For both types of profiles, taxonomic and functional, we use a two-layers Bi-LSTM network, built with Tensorflow-GPU Keras 2.6.0 (see https://www.tensorflow.org/). Output is the prediction of the “theater of activity”.

The first Bi-LSTM layer, with 128 units, is fed by a tensor of shape (117 727, 1), in the case of taxonomy, or of shape (13 041, 1), in the case of function, respectively. Weights are initialized by setting `kernel_initializer` to `glorot_uniform`. The second Bi-LSTM has 64 units. Both layers employ L2 regularization to avoid overfitting.

Both two-layers Bi-LSTM networks are combined using a fully-connected layer with a softmax function for multi-category classification as a final layer. After training the model for 59 epochs using the Adam optimizer, an accuracy of 95.24% was achieved on the test set. Although this is lower than the accuracy of 98.61% achieved using the DeepToA framework described above, it suffices for the purpose of feature importance.

We use SHAP (SHapley Additive exPlanations) values to determine which taxonomic and functional features are important for the prediction (Lundberg and Lee, 2017). In more detail, we use the SHAP deep explainer module (see https://github.com/slundberg/shap) to analyze our model. We randomly sample half of the input samples and used these for training (due to computational constraints). Then SHAP values are computed for all features using the full test set, and the results are summarized in Figure 2 and Figure 3.

**Fig. 2.**
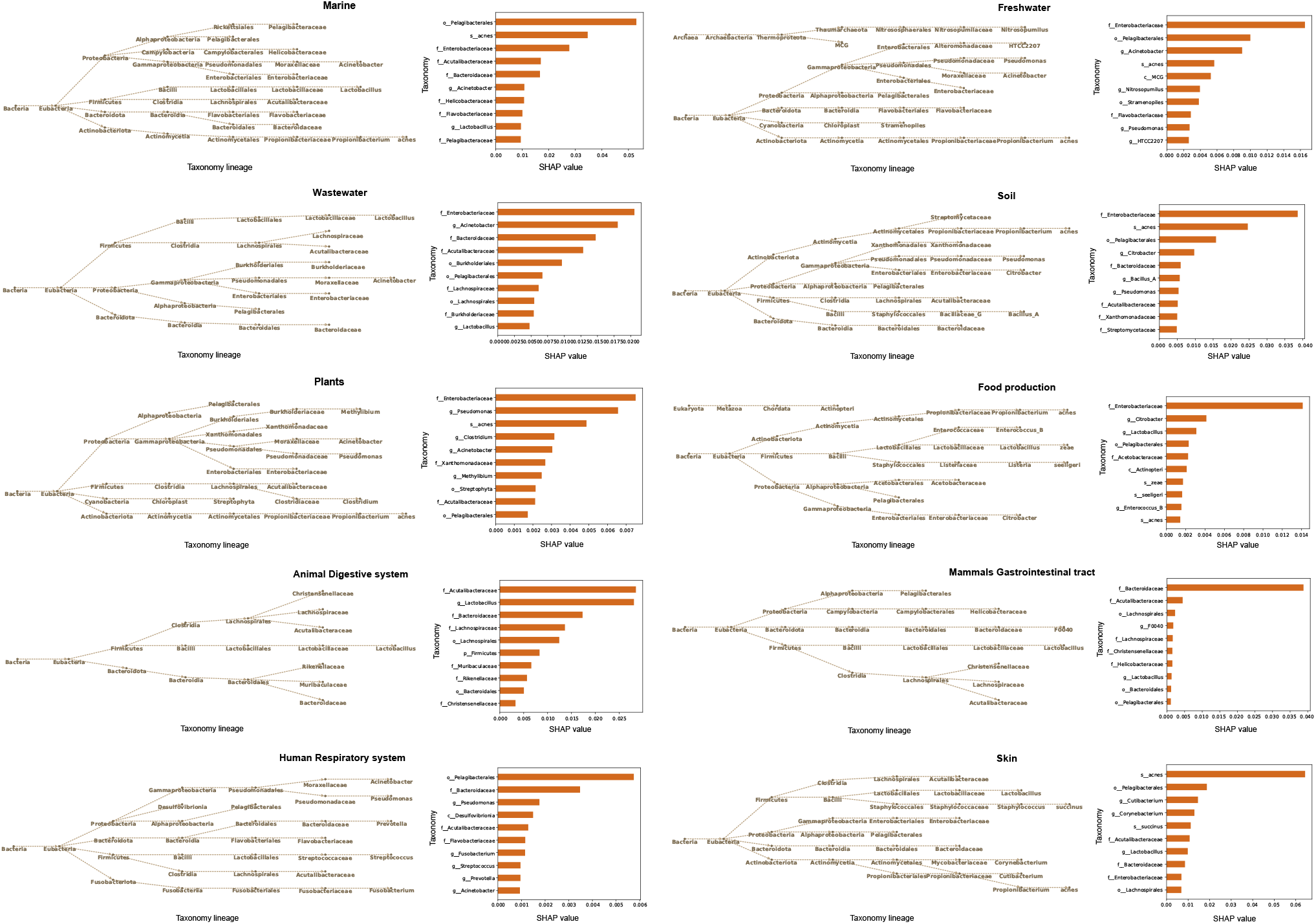
For each of the ten “theaters of activity” under consideration, we show a bar chart of the ten largest SHAP importance values for taxonomic features, together with a display of the corresponding taxonomic lineages.

**Fig. 3.**
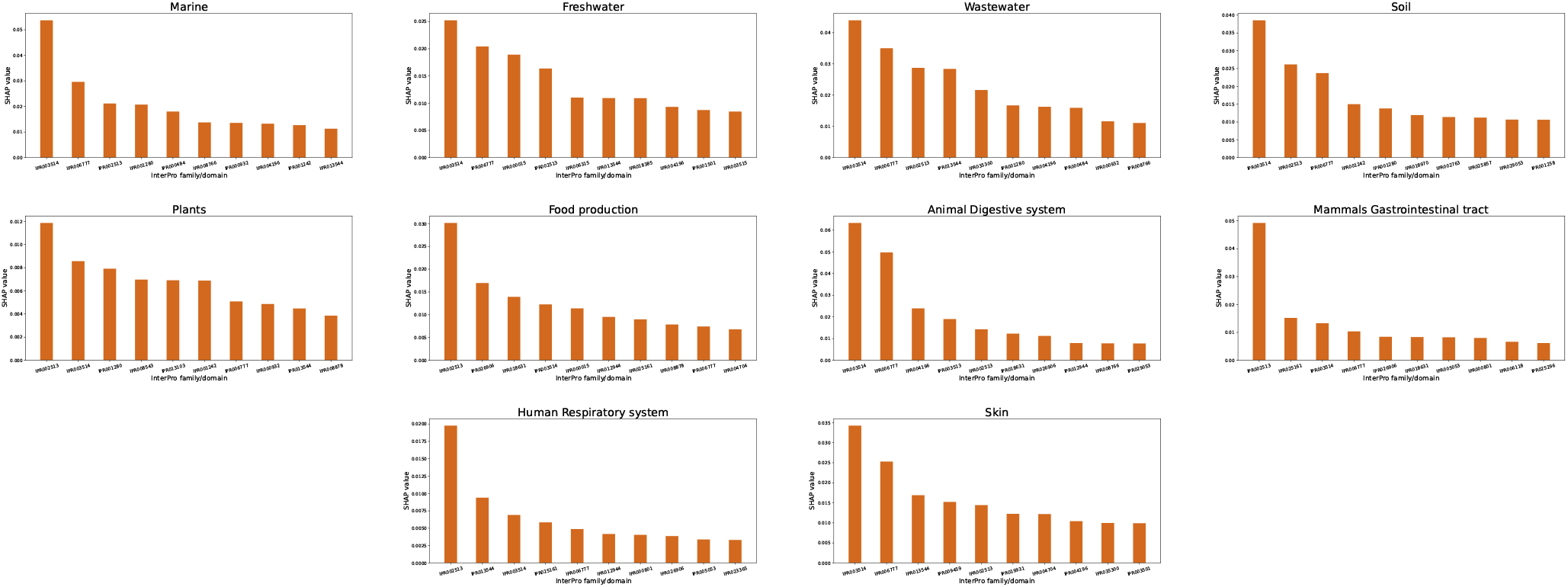
For each of the ten “theaters of activity” under consideration, we show a bar chart of the ten largest SHAP importance values for functional features (InterPro families).

## 3 Result

### 3.1 Performance of DeepToA

To develop an accurate classification model for determining the “theater of activity” for a metagenomic dataset, we explored several ways of combining taxonomic and functional data with different neural network techniques. First, we considered using either the initial taxonomic profile, or the initial functional profile, separately, as input for neural network model. Second, we considered using either of the processed profiles as input. Third we investigated two different approaches to combining both taxonomic and functional data. The accuracy achieved for each of these combinations is reported in Table 2.

**Table 2.**
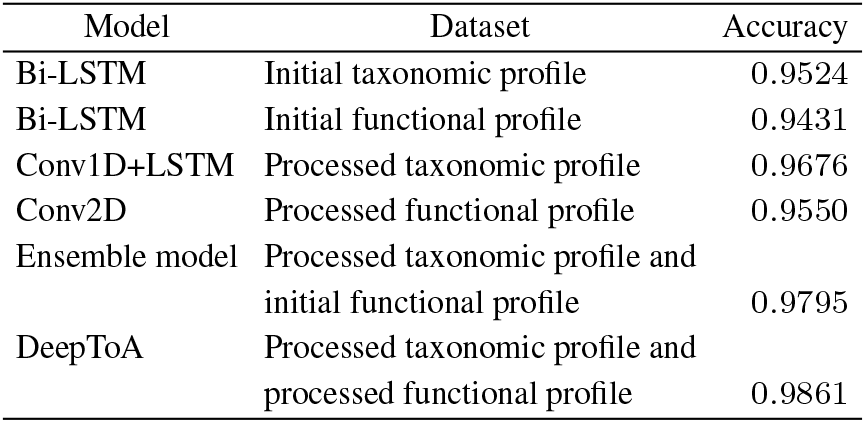
Model evaluation. For different deep-learning models and for different choices of dataset, we report the accuracy of ToA prediction.

The accuracy achieved when using a straightforward model built on the initial taxonomic profiles, or on the initial functional profiles, is 95.24%, or 94.31%, respectively. In comparison to these baseline values, the DeepToA model achieves an accuracy of 98.61%.

### 3.2 Additional information increases precise accuracy

In the third row of Table 2, we report on the performance on processed taxonomic profiles. Here we fed the input data into three one-dimensional CNN layers, each equipped with a ReLU activation function and max-pooling layer. This is followed by two layers of LSTM, each with L2 regularizers with 0.001 and dropout rate 0.1. We use a dense layer with ten units and softmax activation function as the first model’s output layer.

This was then trained using the Adam optimizer with initial learning rate 0.003, decreasing conditionally on the accuracy on the validation set. An accuracy of 96.23% was achieved on the test dataset. This suggests that the processing of taxonomic features captures evolutionary information and this improves prediction ability.

In the fourth row of Table 2, we report the achieved performance on processed functional profiles, using a two-layers CNN, as described above. Because the sample shape is rectangle, the filter size, strides and max-pooling size are set to a rectangle shape. An accuracy of 95.50% was achieved on the test dataset. This indicates that adding external textual information has a positive effect on prediction.

### 3.3 Model result interpretation

To determine which taxonomic and functional features play a major role when classifying the theater of activity of a sample, we built a two-layer Bi-LSTM model on both the taxonomic and functional data, and then applied the Deep SHAP method to obtain SHAP feature importance values.

#### 3.3.1 Feature importance for taxa

In Figure 2, for each of the ten “theaters of activity” (ToA) under investigation, we list the ten taxa that have the highest SHAP values for that particular ToA. In addition, we display the lineage of each such taxa using a part of the taxonomy.

The taxa listed for a particular ToA are often taxa that are known to be be associated with the ToA. For example, the Enterobacteriaceae family shows high importance in freshwater, wastewater, soil, plants and food production. Likewise, the Pelagibacterales order shows high importance for the marine environment and is an order composed of free-living marine bacteria that make up roughly one in three cells at the ocean’s surface (Wikipedia, https://en.wikipedia.org/w/index.php?title=Pelagibacterales, accessed 11-March-2022). However, it is less obvious why it should appear as the most important taxonomic feature for human respiratory system, and also shows up in human skin and food production. Similarly, while *Propionibacterium acnes* has high importance for human skin, it is also listed for marine, freshwater, plants and soil.

#### 3.3.2 Feature importance for function

Functional profiles considered here are based on InterPro families and domains, which are identified by IPR accession numbers. As shown in Figure 3, IPR003514 (Microviridae F protein family) has the highest importance for the prediction of Marine, Freshwater, Wastewater, Soil, Animal Digestive System and Skin. IPR002513 (tract Tn3 transposase DDE domain) has highest importance for the prediction of Plants, Food Production, Mammals Gastrointestinal and Human Respiratory System.

## 4 Discussion and conclusion

Here we introduce DeepToA, an ensemble deep learning framework that aims at predicting the “theater of activity” (ToA) of a microbiome from the taxonomic and functional profiles of its metagenome. To the best of our knowledge, this is one of the first deep-learning approaches to focus on metagenomic data, rather than 16S community profile data, and to utilize both taxonomic and functional profiles (Shenhav *et al*., 2019; Zha *et al*., 2020; Wu *et al*., 2021).

In addition to the ToA classifier, we also provide explanations in terms of both the initial taxonomic and functional profiles. We see that, first, not surprisingly, taxa known to be tightly associated with a particular ToA can have a high associated importance score. However, there are also less obvious appearances, such as *P. acnes* in Marine, and Pelagibacterales in Human Respiratory System and Skin.

We also provide a pre-trained embedding matrix specifically for mapping textual taxonomy information to a numeric vector.

As a machine-learning approach, DeepToA will benefit from increases in the amount of data available for training. With a further increase in the number of sequenced genomes and metagenomes, it will be possible to improve DeepToA so as to distinguish between a larger number of “threaters of activity”, including “cryptic” ones that are not obvious during sample collection. While we focus here on distinguishing between 10 diverse “theaters of activity”, we envision future classifiers addressing a much finer classification, between a “healthy” and “diseased” human respiratory system, say.

## Supporting information

Training data

Test data

## Availability of data and materials

Web server and data is available at http://ab.inf.unituebingen.de/software/deeptoa.

## Authors’ contributions

D.H.H., A.G. and W.Z. designed the study. W.Z. developed the machine learning approach and carried out model based analysis. A.G. and W.Z. performed the microbiome analysis. D.H.H, W.Z. and A.G. and wrote the article. A.G. designed the web server. A.G. and W.Z. packaged the software. All authors discussed the results and edited the manuscript.

## Acknowledgements

We acknowledge hardware support by the High Performance and Cloud Computing Group at the Zentrum für Datenverarbeitung of the University of Tübingen, the state of Baden-Württemberg through bwHPC, the German Research Foundation (DFG) through grant no. INST 37/935-1 FUGG. We also acknowledge support of the BMBF-funded de.NBI Cloud within the German Network for Bioinformatics Infrastructure (de.NBI) (031A532B, 031A533A, 031A533B, 031A534A, 031A535A, 031A537A, 031A537B, 031A537C, 031A537D, 031A538A).

